# A structural and mechanistic model for BSEP dysfunction in PFIC2 cholestatic disease

**DOI:** 10.1101/2024.07.29.605648

**Authors:** Clémence Gruget, Bharat Reddy, Jonathan Moore

**Affiliations:** Massachusetts Institute of Technology, Cambridge, MA, USA; Rectify Pharmaceuticals, Cambridge, MA, USA

## Abstract

BSEP (*ABCB11*) transports bile salts across the canalicular membrane of hepatocytes, where they are incorporated into bile. Biallelic mutations in BSEP can cause Progressive Familial Intrahepatic Cholestasis Type 2 (PFIC2), a rare pediatric disease characterized by hepatic bile acid accumulation leading to hepatotoxicity and, ultimately, liver failure. The most frequently occurring PFIC2 disease-causing mutations are missense mutations, which often display a phenotype with decreased protein expression and impaired maturation and trafficking to the canalicular membrane. To characterize the mutational effects on protein thermodynamic stability, we carried out biophysical characterization of 13 distinct PFIC2-associated variants using in-cell thermal shift (CETSA) measurements. These experiments reveal a cluster of residues localized to the NBD2-ICL2 interface, which exhibit severe destabilization relative to wild-type BSEP. A high-resolution (2.8 Å) cryo-EM structure provides a framework for rationalizing the CETSA results, revealing a novel, NBD2-localized mechanism through which the most severe missense patient mutations drive cholestatic disease.

## Introduction

ABCB11, more commonly known as BSEP (bile salt export pump), is a member of the ABC transporter superfamily, and functions to transport bile salts across the hepatocyte canalicular membrane. Loss of BSEP function results in impaired bile acid flow and retention of bile acids in the liver and is linked with cholestatic diseases such as progressive familial intrahepatic cholestasis type 2 (PFIC2), benign recurrent intra-hepatic cholestasis type 2 (BRIC2) and intrahepatic cholestasis of pregnancy (ICP) ^1-5^.

Of these, PFIC2, with a prevalence of 1/50,000 -1/100,000 people ^6^ is the most severe. PFIC2 arises due to biallelic mutations in BSEP and has an early onset, usually in infancy. PFIC2 is characterized by hepatic bile acid accumulation, which leads to hepatotoxicity resulting in fibrosis, cirrhosis, increased risk of hepatocellular carcinoma, and ultimately, liver failure ^7-11^. The severity and onset of PFIC2 disease is dependent on the degree to which bile acids can be exported from hepatocytes. Nonsense and frameshift mutations give rise to the most severe and earliest onset forms, while missense mutations, depending on their level of dysfunction, drive varying degrees of severity ^12^. Beyond monogenic disorders, emerging evidence links BSEP dysfunction to the pathogenesis of multifactorial liver diseases such as drug-induced liver injury ^13,14^, biliary atresia ^15^, primary intrahepatic stones ^16^, and MASH (formerly NASH) ^17-19^.

At the cellular level, PFIC and BRIC mutations in BSEP have been shown to cause complete loss of expression, decreased mRNA/protein stability, impaired trafficking, or diminished transport ^2,5,20-23^. The most common mutations are missense trafficking mutations ^12^, where most newly synthesized BSEP is recognized by cellular quality control in the ER and targeted for degradation. Several mutations in BSEP occur frequently, including the E297G and D482G variants, present in 58% of European PFIC2 patients ^10^. Reduced cell surface expression of BSEP missense mutants due to defective trafficking is analogous to the most common disease-causing mutations that occur with the cystic fibrosis transmembrane conductance regulator (CFTR). As with CFTR, trafficking defects can be corrected when protein is expressed at low temperature or in the presence of chemical chaperones ^20,22,24,25^, and it is believed that small molecule trafficking modulators, or “correctors”, may be a possible approach to partially restore expression of BSEP at the canalicular membrane of hepatocytes. Several published studies demonstrate that trafficking dysfunction in mutant BSEP may be corrected by small molecules that directly target the mutant protein.

For example, CFTR correctors have been shown in vitro to rescue trafficking and functional defects for PFIC-associated ABCB11 and ABCB4 mutations ^26-28^. The pharmacological chaperone 4-phenylbutyrate (4-PBA) has also demonstrated the capacity to correct protein folding and trafficking of these and other ABC transporter targets ^22,24,29^. In a clinical study, short-term 4-PBA treatment in a pediatric patient with PFIC2 resulted in increased BSEP expression at the canalicular membrane, decreased serum bile acid levels, and lower pruritus scores ^30,31^. These data indicate the potential for developing small-molecule ABCB11 and ABCB4 positive functional modulators for the treatment of PFIC2/3 and other cholestatic diseases.

While it is known that BSEP missense mutations lead to protein trafficking defects and a resulting loss of function, we currently lack an understanding of the mechanisms through which this occurs and, importantly, if these mechanisms can be modulated by drug molecules to achieve a therapeutic benefit. Rigorous, detailed characterization of both the structural and mechanistic effects of disease-causing mutations in BSEP may address this gap in knowledge and, ultimately, inform strategies for the discovery of urgently needed therapies for PFIC2 and other cholestatic diseases.

Protein trafficking defects due to missense mutations arise as a result of protein misfolding. There are several potential pathogenic folding mechanisms that lead to loss of function and the disease state. For example, folding of local structural elements, protein domains, or defective assembly of multidomain proteins may all be affected by missense mutations and can occur either co-translationally or post-translationally. Depending on the cellular environment and specific folding pathways, the mechanisms through which misfolded proteins are recognized by cellular quality control and targeted for degradation can vary. Many studies have demonstrated that protein folding defects often lead to a decrease in the thermodynamic stability of the nascently or fully folded protein ^32^, and may be observed directly through the experimental determination of the melting temperature of the target protein ^33,34^. More recently, methods for the determination of melting temperature in a cellular environment have been developed ^35-37^, and possess the advantage of characterizing the energetics of folding in the context of the full proteostasis network, including interactions with chaperones or accessory proteins that may be required for proper folding, cellular localization and function.

To characterize the effects of BSEP disease-causing mutations on protein thermodynamic stability, we have carried out in-cell thermal shift (CETSA) measurements for 13 different patient BSEP mutations, including the most prevalent variants, E297G and D482G. Using a novel split luciferase detection method, shifts in aggregation temperature (T_agg_) could be binned into three groups: a) mutations with no effects on T_agg_, b) mutations causing mildly destabilizing thermal shifts of -2-4°C, and c) mutations causing severe destabilization of 9-11°C. All highly destabilizing mutations were localized to the NBD2-ICL2 interface of BSEP. Confocal microscopy indicated cytoplasmic versus plasma membrane (PM) localization in HEK293 cells, confirming that these mutations resulted in defective protein trafficking. To determine the structural basis for BSEP protein destabilization, we determined the cryo-EM structure of wild-type (WT) BSEP to 2.8 Å resolution. Focusing on mutations in NBD and ICL2, our high-resolution cryo-EM model provides a structural framework for rationalizing the thermal destabilization of these mutants, suggesting a novel, NBD2-localized mechanism through which the most severe missense patient mutations drive severe cholestatic disease.

## Results

### Thermal stability of BSEP patient variants

BSEP variants drive PFIC2 disease through decreased BSEP expression and function. Our study aimed to determine if protein thermodynamic stability is linked to protein trafficking defects and diminished function associated with previously characterized biallelic BSEP variants ^12,20^. We initially focused on the missense mutations E297G and D482G, as these are the most common patient variants, occurring in 58% of European patients ^10^. Based on the results from these initial experiments, we then expanded our characterization to other known patient variants localized to the BSEP nucleotide binding domains (NBDs) and intracellular loops that interact with NBDs. These mutations range from benign variants, including several BSEP polymorphisms, to mutations that result in mild disease and severe cholestatic disease. The variants we examined are listed in Table 1 and highlighted in our high-resolution cryo-EM model of BSEP in Figure 1.

**Table 1.**
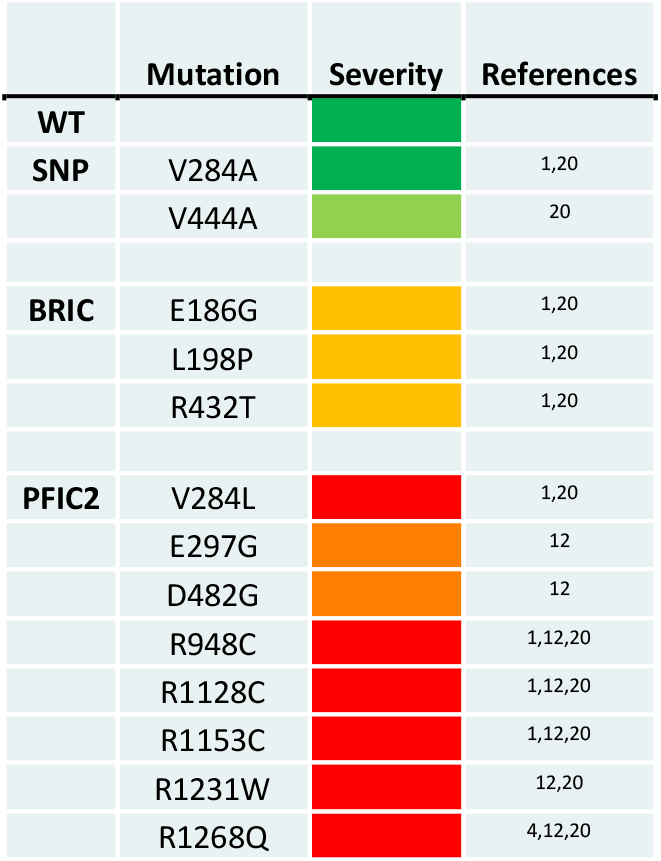
BSEP variants selected for this study. WT and SNPs, BRIC, and PFIC2 variants, are color coded in green, light green, amber, orange, and red to reflect disease severity.

**Figure 1.**
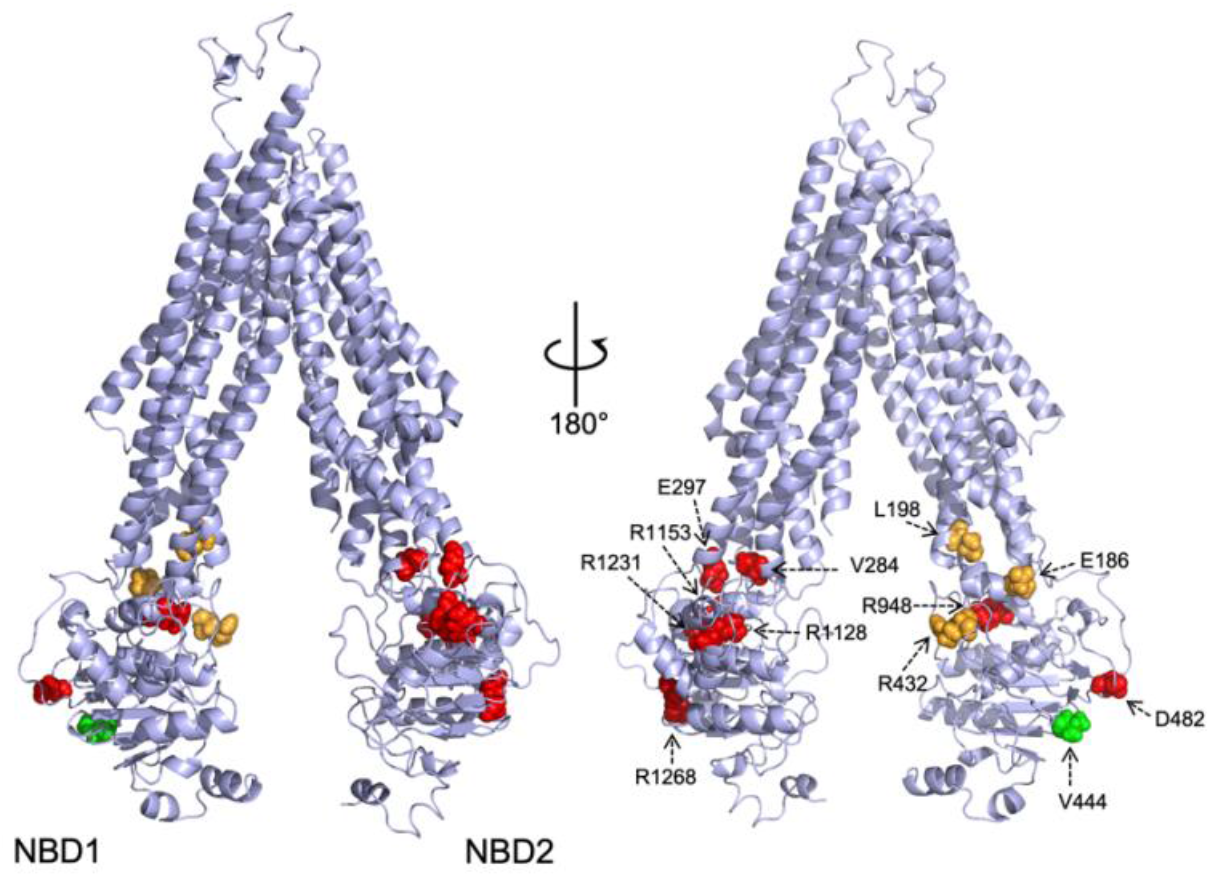
Variant residues are highlighted in a high-resolution structural model of WT BSEP. Sidechain color reflects the disease severity from Table 1. All variants chosen are in the NBDs or ICLs.

To examine the thermal stability of proteins in a cellular environment, we carried out CETSA experiments using a live cell HiBiT split luciferase detection (Promega, Madison, WI)) of non-aggregated BSEP and BSEP variants. Using a strategy like that described in the literature ^38-41^ (Figure 2), we developed plasmid constructs containing the BSEP protein, followed by a C-terminal GSGS linker and an 11-amino acid HiBiT tag. Briefly, plasmids containing the appropriate BSEP construct were transfected into HEK293 cells for 24 hours. Cells were harvested and transferred to PCR tubes, heated in a PCR block at the appropriate temperature for 3 minutes. Cells were then lysed, and luciferase activity was reconstituted by addition of the LgBiT nanoluciferase fragment and furimazine substrate, followed by transfer to a 384-well plate for measurement of luminescence in a plate reader.

**Figure 2.**
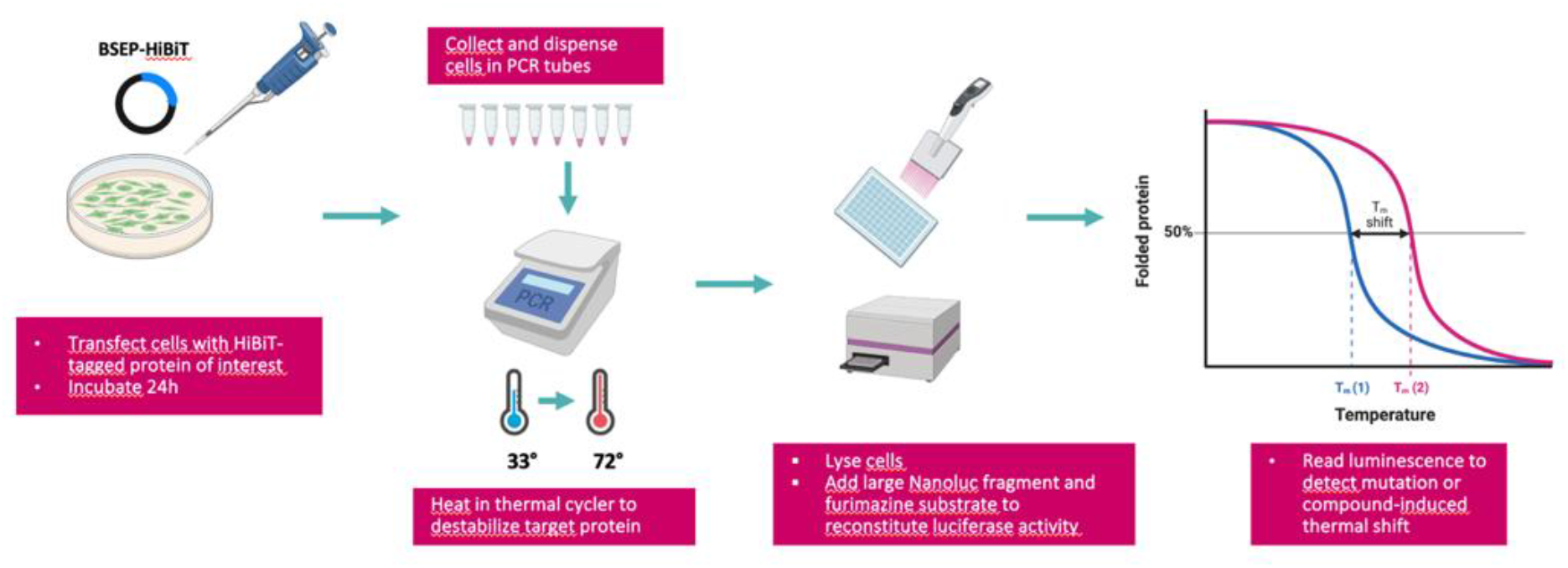
Schematic overview of a luminescence based cellular thermal shift assay (CETSA) workflow. HEK293T cells transfected with BSEP constructs with a C-terminal HiBiT tag are thermally challenged before being lysed and the Large NanoLuc fragment (LgBiT) and substrate are added to generate a luminescence signal. A melting/aggregation temperature is extracted from the resulting melting curves to detect a potential thermal destabilization of BSEP mutations/SNPs

We first aimed to characterize four BSEP proteins: wild type, the common polymorph V444A, and the most common, but mild disease mutations E297G and D482 (Figure 3).

**Figure 3.**
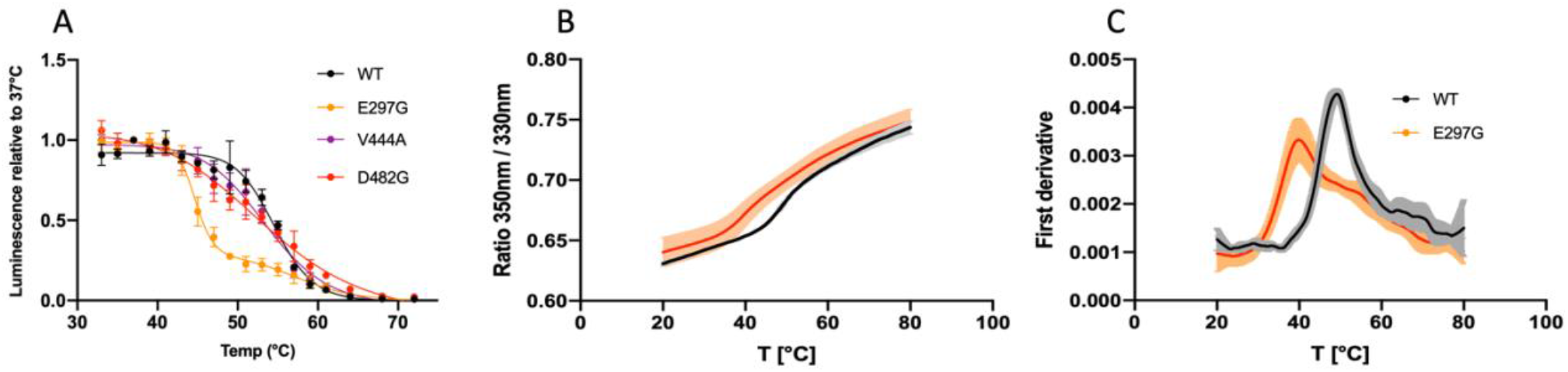
(A) Thermal stability curves of wild-type BSEP and variants E297G, D482G and the SNP V444A. Each point represents mean ± SD of at least three independent experiments. Curves were fitted with a non-linear regression model in GraphPad Prism (Version 9, Boston, MA) and an aggregation temperature (T_agg_) was obtained. (B-C) Melting profile of purified wild-type BSEP and E297G mutant obtained by nanoDSF. B: 350/330 fluorescence emission. C: First derivative of the 350/330 ratio signal.

CETSA curves (Figure 3A) for WT and V444A exhibit nearly the same aggregation temperatures (T_agg_) of 54.46 °C and 53.38 °C, which is expected as V444A homozygotes are healthy, do not display a disease phenotype, but may be more susceptible to cholestatic conditions such as drug-induced liver injury (DILI) and intrahepatic cholestasis of pregnancy (ICP) ^13,42-44^. The D482G BSEP variant shows a similar T_agg_ to that of V444A and WT BSEP within experimental error. So, while this variant has been previously characterized as a low protein-expressing patient variant, the low expression observed in this and other studies does not arise due to destabilization. In contrast to WT, V444A and D482G BSEP variants, E297G BSEP displayed a very high degree of destabilization, with a T_agg_ of 44.63°C, a shift of ∼9.8°C.

To verify whether the aggregation temperatures (T_agg_) observed for WT and E297G BSEP in live cell CETSA accurately reflect the melting temperatures of these proteins when purified, we conducted nanoDSF experiments on BSEP proteins that had been biochemically purified to homogeneity. The results, shown in Figures 3B and 3C, show melting temperatures (T_m_) for WT and E297G BSEP of 48.96 ± 0.45°C and 40.23 ± 0.38°C, respectively, yielding a ΔT_m_ of 8.73°C. This difference is slightly less than the 9.83°C ΔT_agg_ observed in the CETSA experiments. Although direct quantitative comparisons between CETSA and nanoDSF transition temperatures are difficult, the large ΔT_m_ and ΔT_agg_ values indicate similar relative stabilities for BSEP proteins in cellular and purified states. Therefore, relative protein stability is not significantly affected by the cellular environment.

As the magnitude of destabilization for E297G BSEP was unexpected, we broadened our study to examine additional mutations in the NBDs of BSEP. Based on a literature survey, an additional set of patient variants representing a range of potential disease severities was chosen. Utilizing the highly sensitive HiBiT split luciferase detection system enabled us to assess the thermal stability of mutants that previously showed very low or undetectable expression levels ^20^. Complete CETSA curves for WT BSEP and 13 reported patient variants are shown in Figure 4.

**Figure 4.**
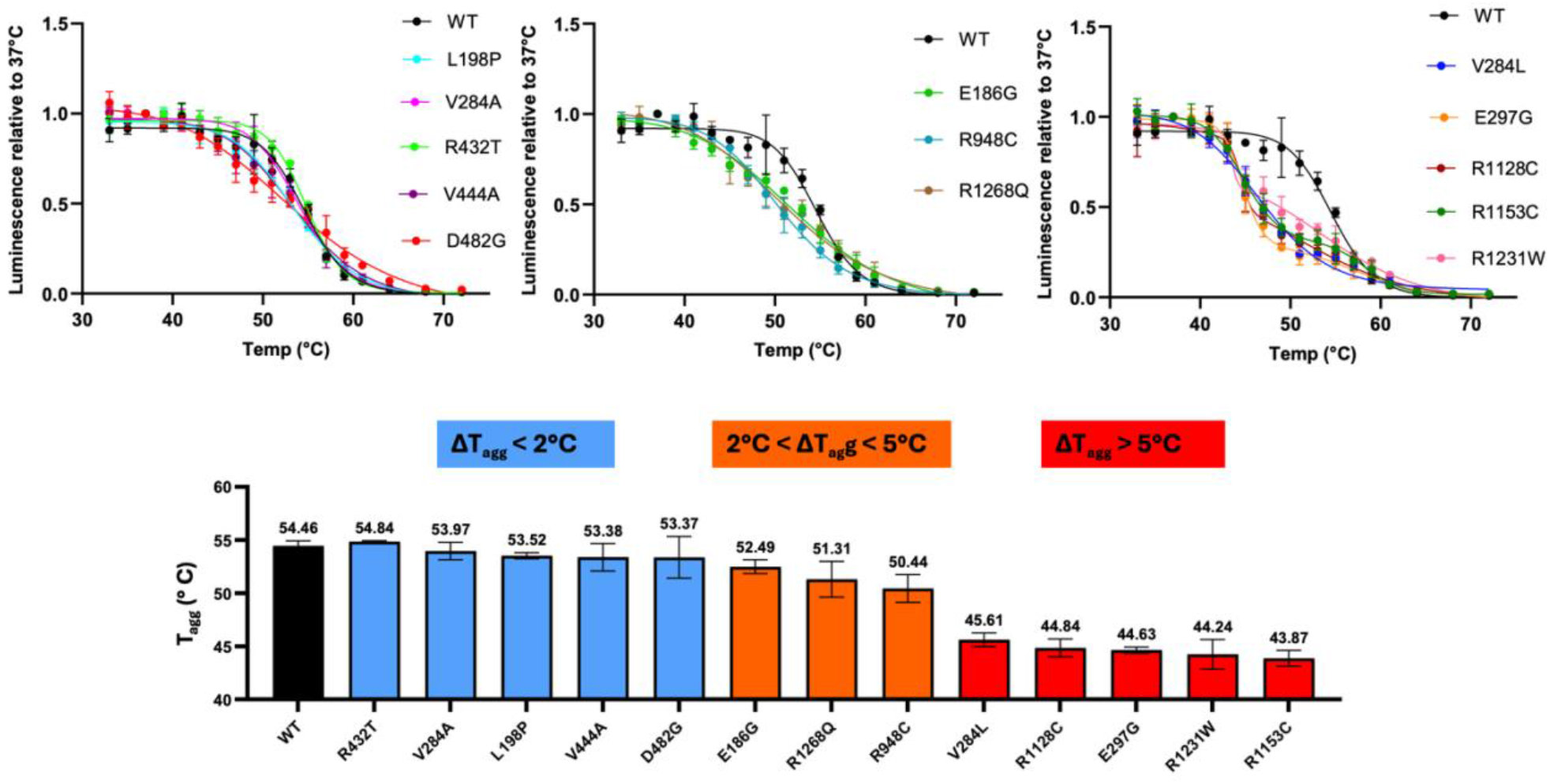
(Top) Thermal stability curves of wild-type BSEP and thirteen patient variants. Each point represents mean ± SD of at least three independent experiments. Curves were fitted with a non-linear regression model in GraphPad Prism (Version 9, Boston, MA) and a T_agg_ value was obtained. (Bottom) T_agg_ values derived from the curves in A), shown from highest to lowest. ΔT_agg_ represents the difference in aggregation temperature induced by missense mutations relative to wild type. Each value corresponds to the mean ± SD T_agg_ obtained from at least three independent melting curves fitted individually.

Here, we have binned the BSEP variants into three groups. One group, corresponding to WT BSEP, the V444A and V284A polymorphs, and the BRIC variants L198P, R432T show either minimal (< 2°C) or no destabilization. The mild BSEP disease mutation D482G also falls into this category, showing only a 1.1 °C ΔT_agg_ compared to WT BSEP. Another group of variants, consisting of the BRIC variant E186G, and PFIC2 variants R948C and R1268Q, exhibit an intermediate range of destabilization with ΔT_agg_ between 2-5°C, with both PFIC2 variants showing ΔT_agg_ values in the range of 3-4°C. The third group of variants, all representing either mild (E297G) or severe (V284L, R1128C, R1153C, R1231W) PFIC2 patient mutations, show a very high degree of destabilization, with ΔT_agg_ values ranging from 8.8-10.6 °C. Notably, all these highly destabilized variants are localized to the NBD2 domain of BSEP, suggesting that this domain may represent a hotspot for destabilization and could play a critical role in BSEP folding.

### Cellular localization of BSEP variants

As all the BRIC and PFIC2 variants we examined have been previously reported to be low expressing or protein trafficking mutants, we next characterized the cellular localization of BSEP WT and variants using Western blots and confocal microscopy. We hypothesized that if CETSA destabilization correlates with protein misfolding, variants with lower aggregation temperatures (T_agg_) would be more susceptible to degradation via endoplasmic reticulum-associated degradation (ERAD). This susceptibility would be directly reflected by the distribution of the ER and PM protein forms in Western blots, as well as through direct observation of ER and PM markers in confocal images.

ABC transporters such as CFTR and BSEP typically possess two distinct bands in Western blots. The lower molecular weight, or B-band, represents core glycosylated, ER-localized protein, while protein that reaches the Golgi becomes complex glycosylated, higher molecular weight C-band protein that may be either plasma membrane (PM) or intracellularly localized. Trafficking-deficient variants usually show decreased protein expression relative to wild-type, increased B-band protein, and diminished or no C-band protein. As expression levels may vary in plasmid-based experiments using transient transfection, only a qualitative assessment of the Western blots is possible. The Western blot for WT BSEP (Figure 5D) shows an intense C-band and a much weaker B-band, indicative of fully glycosylated BSEP protein with little ER-localized protein. R432T BSEP appears to have a very similar C/B band ratio as WT, while for V284A and V444A, we also observe intense C-bands, but a slightly lower C/B ratio. These ratios align well with the merged images in Figure 5A, which show predominant staining at the plasma membrane, with trace amounts of protein in the ER. Variants that show both ER and PM localization by Western (E186G, L198P, V284L, E297G) (Figure 5B) also show both ER and PM staining in the confocal images. Of these, the BRIC variants E186G and L198P, and the common PFIC2 variant E297G all show low expression with protein equally distributed between B and C bands in the Westerns, and the ER and PM in the confocal images, while V284L clearly shows more B-band protein in the Western blot, and a slightly brighter ER component via immunofluorescence (IF). Finally, the variants D482G, R948C, R1128C, R1153C, R1231W, and R1268Q all localize predominantly in the ER, with little to no cell surface localization, which is confirmed by the merged confocal images in Figure 5C.

**Figure 5.**
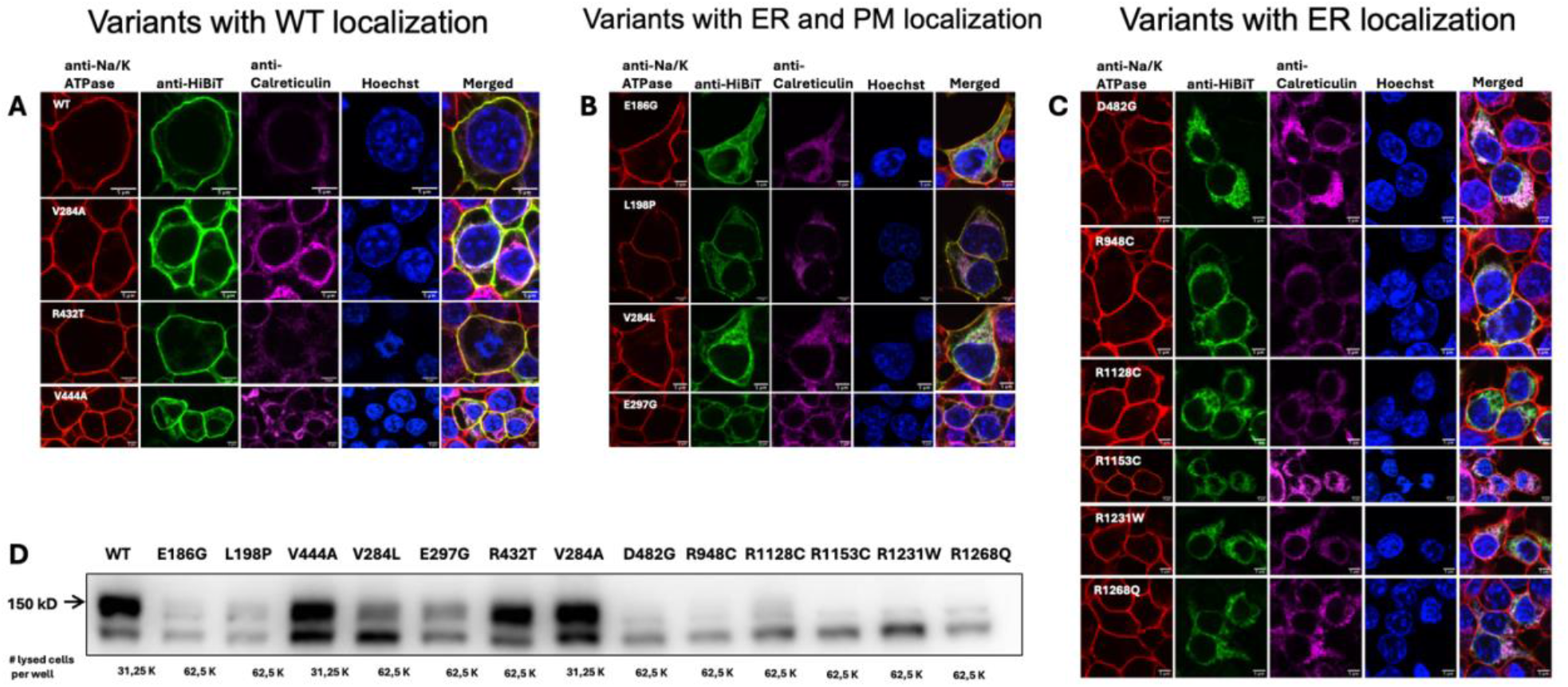
Cellular localization of BSEP wild-type and variants. A-C: Immunofluorescence confocal images of HEK293T cells expressing C-terminal HiBiT-tagged BSEP constructs. Red, green, magenta, and blue fluorescence correspond to PM, BSEP protein, ER and nuclear markers, respectively. Scale bars are 5 μm. A) Variants with WT PM localization, B) BSEP variants localized to both the PM and ER; C) BSEP variants found primarily in the ER. D) Western-Blot of HEK293T cell lysates transfected with BSEP wild-type and mutants containing a C-terminal HiBiT tag. The top band (∼150 kD) represents the mature, fully glycosylated form of BSEP while the bottom band represents an immature, core glycosylated form of BSEP.

While most expression and localization data correlate well with protein stability, one clear outlier is the D482G variant. D482G BSEP showed no protein destabilization within experimental error in the CETSA measurements yet showed little mature protein in Western blots and almost complete ER localization in the confocal images, suggesting that additional factors beyond thermodynamic stability are likely at play. These may include alternative aspects of cellular trafficking or processing that are yet to be fully understood. An earlier study has proposed alternative gene splicing as a driver for disease pathology for D482G ^20^, but this can be ruled out in the present study given the nature of the *in vitro* methods employed.

Another mutation, R432T, which has been characterized as a BRIC mutation with slightly reduced expression levels in previous studies, ^20^ appears to display near wild-type maturation and localization in Western blots and confocal microscopy. This suggests other functional impairments, such as ATP binding or defective transport activity, as a basis for pathogenicity. Recent structures of the BSEP in the ATP-bound state support the notion that R432 may be important for proper NBD dimerization and transport function rather than affecting protein folding and assembly ^45^.

### Structural basis of destabilization for BSEP variants

To better understand the structural rationale for the protein destabilization observed for BSEP variants, especially the most highly destabilized, NBD2-localized variants, we solved the cryo-EM structure of BSEP in the inward-facing, open conformation. Our cryo-EM structure is of higher resolution than most previously reported BSEP structures and is particularly well-resolved in the NBDs, where ABC transporters in the inward-facing state typically exhibit increased mobility and consequently decreased resolution compared to the transmembrane domains (TMDs).

Structural models for the five highly destabilized residues identified in the CETSA experiments are shown in Figure 6. Sidechain conformations for all residues discussed are explicitly determined by the cryo-EM density map. Both Val-284 and Glu-297 are in intracellular loop 2 (ICL2). IC2 is formed by the approximately parallel helices 4 and 5 and intracellular helix 2 (IH2) ^45^ and makes close contact with NBD2 (Figure 6A). ICLs in the ABC transporter superfamily play an important role in transducing the dynamic motion of the NBDs due to substrate binding and ATP hydrolysis into structural rearrangement of the TMDs from an inward-facing, closed conformation, to an outward-facing, open conformation. Val-284 is in helix 4, and the cryo-EM model suggests that mutation of Val-284 to a leucine would add steric bulk to the region where helices 4 and 5 pack together, possibly disrupting sidechain electrostatic interactions and packing of E300 and R303. It is not surprising that this mutation would be destabilizing and potentially influence protein folding and domain assembly between TMD2 and NBD2. Glu-297 is located at the base of helix 5, adjacent to IH2. Glu-297 makes several sidechain-backbone H-bonds with Gly-295 and Gly-296 and has one long-range electrostatic interaction with Arg-1153 (Figure 6B) that would be lost in the cysteine variant. Arg-1153 (Figure 6C) is positioned centrally between Glu-297 and Arg-1128. The Arg-1153 sidechain interacts with both the sidechain of Glu-297 and the backbone carbonyls of both Ala-292 and Arg-1128. The sidechain of Arg-1128 (Figure 6D) forms a further electrostatic network with the sidechain of Asp-1131 and the backbone carbonyls of Ser-1144 and Val-1147. Thus, the network of Glu-297, Arg-1153 and Arg-1128 tie together five distinct structural elements within NBD2. R1231, mutated into a tryptophan in PFIC2 patients, is located on the opposite side of IH2 from Arg-1153. The sidechain of Arg-1231 forms a hydrogen bond with the backbone carbonyl of Val-1164 (Figure 6E), acting to stabilize helix 1 of the RecA binding core with the sequence connecting the ABCα domain with the first β-strand of the four-stranded β-sheet. Loss of such an important interaction could significantly impair NBD2 folding.

**Figure 6.**
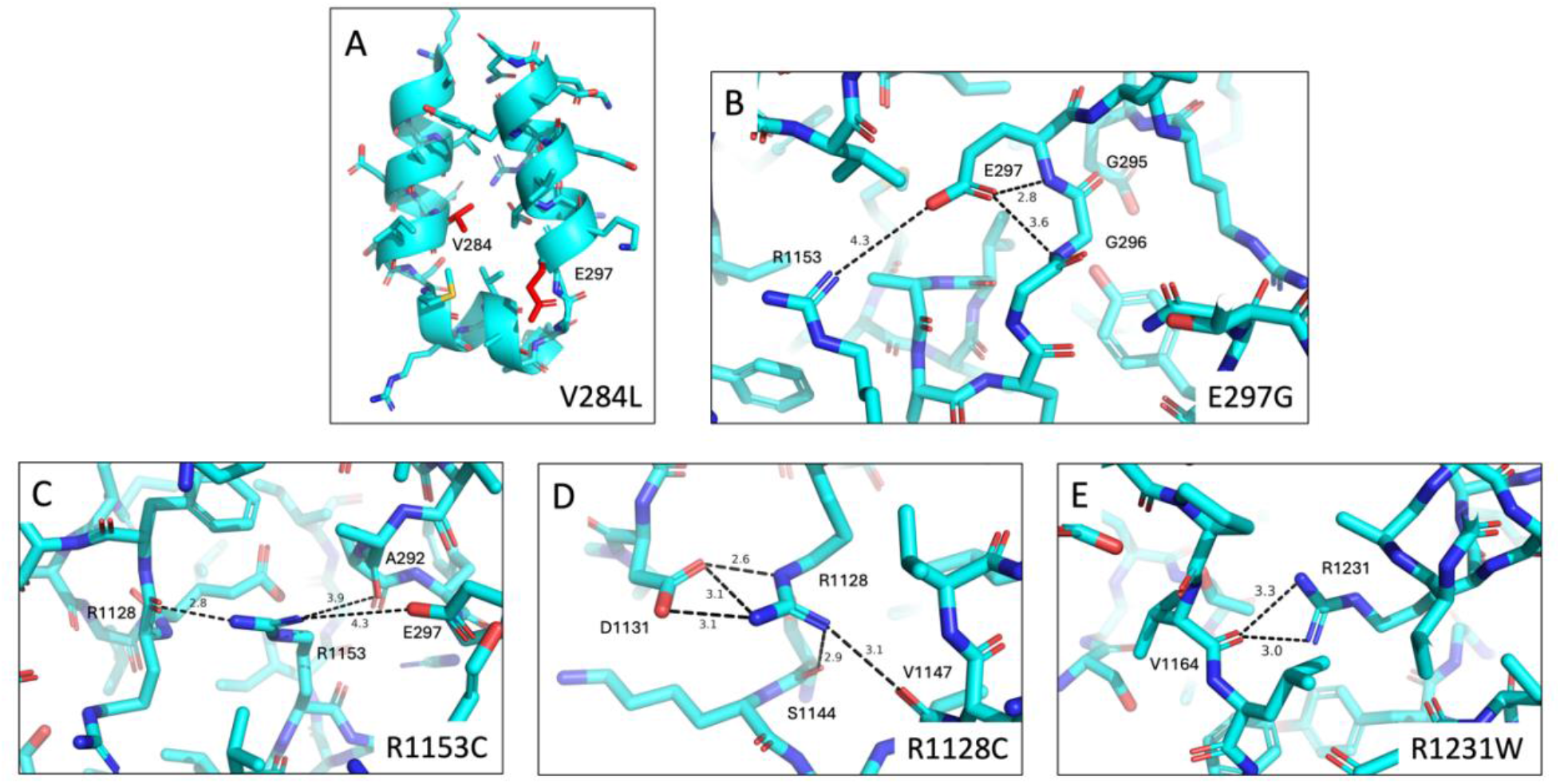
Cryo-EM structural models for five highly destabilized BSEP residues identified from the CETSA experiments. The cryo-EM model was determined to 2.78Å using 435,418 particles for the final reconstruction.

**Figure 7.**
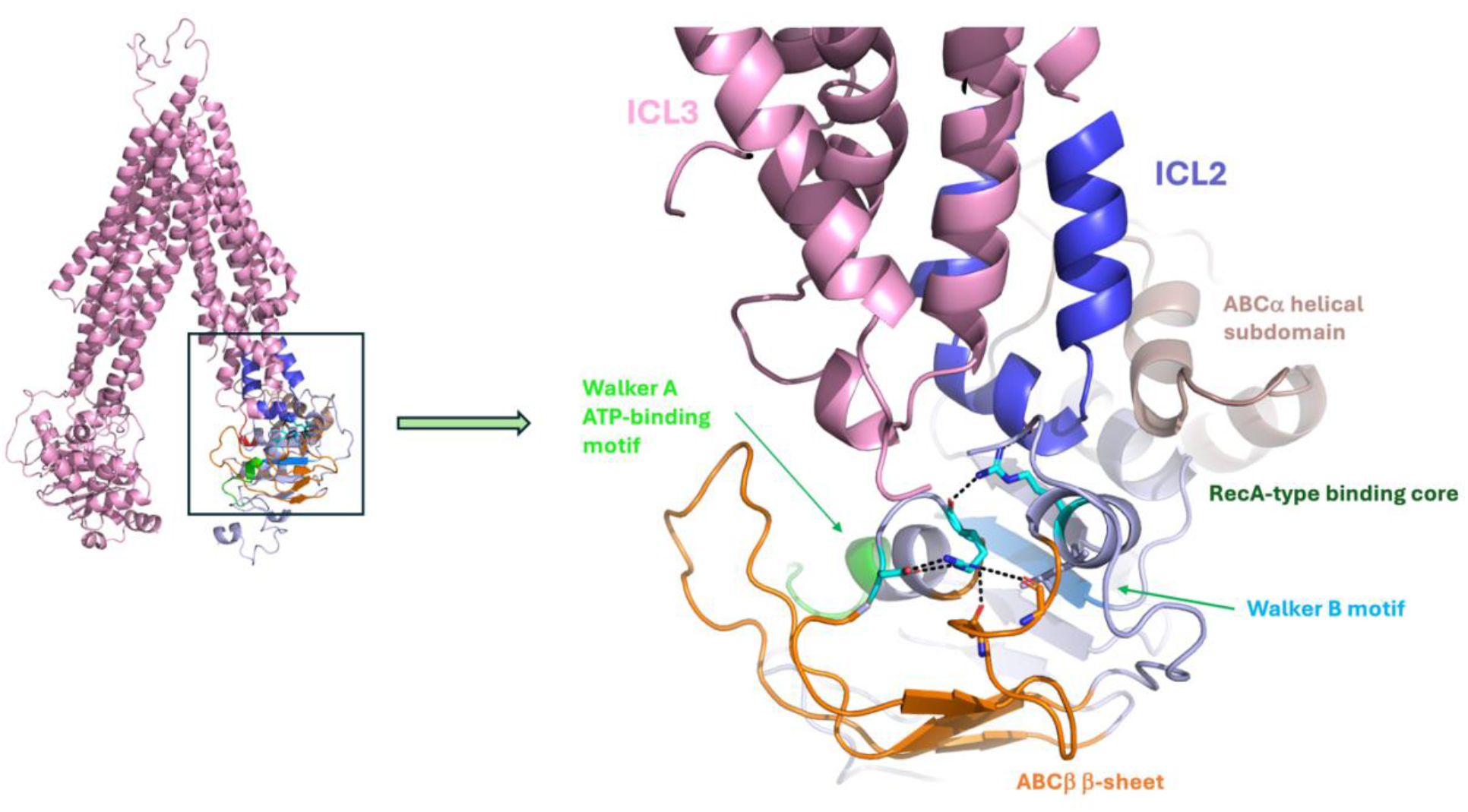
R1128 and R1153 (cyan sticks) form the core of an electrostatic network that stabilizes the NBD2 fold and NBD2-ICL2 interaction by tying together key subdomains of NBD2 such as the Walker A ATP-binding motif, the ABCβ subdomain, helix 1 of the RecA-type binding core, and ICL2 of transmembrane domain 2 (TMD2).

## Discussion

Although it is established that BSEP missense mutations can lead to reduced protein expression and function, the precise molecular mechanisms underlying these effects remain poorly understood. This study provides, for the first time, detailed insights into the molecular basis of protein misfolding induced by BSEP mutations, which contribute to cholestatic diseases such as PFIC2 and BRIC.

Through biophysical, immunofluorescence, and structural analyses, we have demonstrated a clear correlation between the thermodynamic stability of BSEP variants, their cellular localization, and the severity of associated diseases. Notably, our CETSA experiments have identified significant destabilization at the NBD2-ICL2 interface—a critical region for protein folding and function.

While the present study is not an exhaustive analysis of all possible structural interactions, our high-resolution cryo-EM structural data provide a robust framework for understanding the observed protein destabilization. These findings reveal that essential interactions, which maintain the structural integrity of multiple elements within NBD2, are disrupted in many disease-associated variants.

The discovery of hotspots for destabilization within ABC transporters supports the notion that certain protein classes are, by nature, poised at the edge of stabilization ^46^. These proteins require low energy barriers for transitions between conformational states to function effectively, which, for ABC transporters, involves interconversion between inward and outward-facing conformations to facilitate the transport of substrates across biological membranes. While necessary for function, this inherent instability may also make these regions susceptible to destabilizing mutations that can lead to disease.

Given the potential similarities in substrate transport mechanisms across the ABC transporter family, future studies should explore whether the mutation-sensitive structural elements identified in BSEP are conserved in other transporters and how these might relate to pathogenicity in other ABC transporter-mediated diseases.

This study reinforces the notion that protein folding disorders may develop from localized destabilization and misfolding, leading to their recognition and targeted degradation by the cellular proteostasis network. This understanding forms the basis of drug discovery strategies to identify small molecule stabilizers that act as trafficking correctors. Such strategies have previously facilitated the development of transformative therapies for cystic fibrosis ^47^. Our results indicate that designing NBD2-targeted, mechanism-based assays to discover small molecule stabilizers would present a novel and promising approach for correcting BSEP trafficking defects and advancing the development of vital therapies for PFIC2 and other cholestatic diseases.

## Materials and methods

### DNA constructs

Full-length human BSEP cDNA was cloned in a pcDNA vector purchased from Azenta and codon optimized for mammalian cell expression. All point mutations were inserted by inverse PCR. PCR products were ligated and transformed in MACH1 E. coli for clone selection and amplification. All constructs were sequenced by Azenta.

### Cell cultures and transfections

Adherent HEK283T cells were maintained in a humidified atmosphere at 37°C with 5% CO2 in Dulbecco’s modified Eagle’s medium (DMEM) with 4,5 g/L glucose and sodium pyruvate, (Corning) supplemented with 10% fetal bovine serum, 1X GlutaMAX™ and Penicillin-Streptomycin (Gibco).

For CETSA and Western Blot experiments, HEK293T cells were transfected at 70% confluency in a 6-well plate or 12-well plate using PEI MAX^®^ - Transfection Grade Linear Polyethyleneimine Hydrochloride (MW 40,000) (Polysciences) at a 1:3 DNA: PEI mass ratio and harvested 24h after transfection.

For immunofluorescence experiments, HEK293T cells were plated on glass coverslips coated with poly-D-lysine in 24 wells plates and transfected the next day with BSEP-HiBiT plasmid with FuGENE^®^ HD Transfection Reagent according to the manufacturer’s instructions (Promega).

For BSEP protein purification, HEK293S GnT1-cells were grown in suspension in an orbital shaker in FreeStyle™ 293 Expression Medium (Thermo Fisher Scientific) supplemented with 2% fetal bovine serum (Gibco) in a humidified atmosphere at 37°C with 8% CO2. Human BSEP WT and E297G mutant constructs with a C-terminal GFP were expressed in HEK293s GnTI– cells using the BacMam system as described ^48^. Briefly, cells were transduced with baculovirus when cell density reached ∼ 2×106 cells/ml. 8 to 24 hours after transduction, 1mM sodium butyrate was added to the culture, and the temperature shifted to 30°C. Cells were harvested ∼ 72h after transduction, and the pellet was stored at -80°C.

### Western Blots

Adherent HEK293T cells were seeded in a 12-wells plate and transfected at 70% confluency using PEI MAX^®^ - Transfection Grade Linear Polyethyleneimine Hydrochloride (MW 40,000) (Polysciences) at a 1:3 DNA: PEI mass ratio. Cells were harvested 24h after transfection, counted, washed 1x in DPBS, and aliquoted. Cell pellets were stored at -80°C. Cell pellets were resuspended in lysis buffer (0,2% Triton-X in DPBS supplemented with 1X cOmplete™, Mini, EDTA-free Protease Inhibitor Cocktail (Roche)), left on ice for 30 minutes, and spun down 12 000 rpm at 4°C for 10 minutes. The supernatant was collected, and protein separated by SDS-PAGE on 7.5% gel run at 70V for 30 minutes and 80V for 1 hour, transferred on a PVDF membrane using Trans-Blot Turbo transfer kit (BioRad) that was blocked for 2 hours in 5% milk TBST before being probed with (Promega) (CS2006A01 clone 30E5) monoclonal antibody overnight at 4°C (1:4000 in 3%BSA TBST). The next day, the membrane was washed in TBST and incubated for 1 hour with HRP-conjugated goat anti-mouse IgG (sc-2055, Santa Cruz Biotechnology) (1:5000 in 3%BSA TBST). After a final wash in TBST, bands were revealed with SuperSignal™ West Pico PLUS Chemiluminescent Substrate (Thermo Fisher Scientific), and the membrane was imaged on an Amersham Imager 680. To confirm the glycosylation state of B and C-band BSEP proteins, lysates were treated with PNGase F or EndoH (New England Biolabs) in denaturing conditions following the provider’s instructions (Supplementary Figure 1). For better visualization, the amount of cell lysate loaded on gel was twofold for mutants compared with WT.

### CETSA

Adherent HEK293T cells were grown in a 6-wells plate and transfected with BSEP-HiBiT constructs when they reached 70% confluency using PEI MAX^®^ - Transfection Grade Linear Polyethyleneimine Hydrochloride (MW 40,000) (Polysciences) at a 1:3 DNA: PEI mass ratio. Cells were collected 24h after transfection by trypsinization, resuspended in fresh DMEM at a concentration of 1,2.106 cells/ml, and dispensed in PCR tubes (20 μl per tube, 2 replicates per condition). Tubes were placed in a multi-block thermal cycler and submitted to a thermal cycle of 10 minutes pre-incubation at 37°C, 3.5 minutes heating step, and fast cooling to 4°C. Cells were then lysed, and a luminescent signal was generated by adding Nano-Glo^®^ HiBiT Lytic Detection mix prepared per manufacturer’s instructions (Promega) to the heated samples at a 1:1 vol/vol ratio. Cell lysates were subsequently dispensed in a white 384 wells plate (Corning) and luminescence was detected on a plate reader (TECAN Spark^®^).

### Melting curve analysis

For each melting curve, two wells per temperature were averaged. The resulting melting curve was fitted with a non-linear regression model with GraphPad Prism (Version 9, Boston, MA) from which a melting/aggregation temperature was extracted.

### Protein purification

A pellet of cells expressing WT BSEP or E297G mutant was resuspended in solubilization buffer (Tris 50 mM pH8, 150 mM NaCl, 10% glycerol, 2 mM DTT, 2 mM MgCl2, 1%DDM, 0.2% CHS, protease inhibitor tablet and DNase/RNase) for 2 hours at 4°C. The cell lysate was centrifuged at 200000 g for 30 minutes at 4°C, and the supernatant was added to anti-GFP nanobody beads (Protein and Crystallography Facility, University of Iowa) for 1 hour at 4°C. Beads were washed with buffer A (150 mM NaCl, 50 mM Tris pH8, 2 mM DTT, 2 mM MgCl2, 0.02% GDN) and incubated with 3C protease (40 μM) for 3 hours at 4°C to remove the C-terminal GFP tag. The eluted protein was further purified by gel filtration chromatography using a Superose^®^ 6 Increase 10/300 GL (Cytiva) equilibrated with Buffer A.

### Differential scanning fluorimetry

Freshly purified proteins were concentrated to ∼0,4 mg/ml on a Pierce 100K MWCO concentrator (Thermo Fisher Scientific) catalog number 88503 for thermal stability analysis with Prometheus NT48 (NanoTemper Technologies). Tryptophan fluorescence 330/350 nm ratio was measured in high sensitivity capillaries submitted to a temperature ramp of 1°C per minute, from 20 to 80 °C.

### Immunofluorescence and confocal microscopy

HEK293T cells were plated on glass coverslips coated with poly-D-lysine in 24 wells plates. The next day cells were transfected with BSEP-HiBiT plasmid with FuGENE^®^ HD Transfection Reagent according to the manufacturer’s instructions (Promega). 1-2 days after transfection cells were washed three times with cold DPBS, fixed in 100% methanol for 5 minutes, washed again three times with DPBS, permeabilized with 0,1% Triton X-100 for 5 minutes and washed one more time in DPBS. Cells were then incubated in blocking buffer (2%BSA in HBSS) for 30 minutes. Primary antibodies were diluted in the same buffer and added to the cells ON at 4°C (mouse anti-HiBiT (Promega) catalog number CS2006A01 clone 30E5) monoclonal antibody, 1:500; rabbit Alexa Fluor^®^ 647 anti-Calreticulin (Abcam catalog number ab196159), 1:400; rat anti Na/K ATPase (Abcam catalog number ab283345), 1:400). Cells were washed the next day three times in DPBS and incubated 1h RT with secondary antibodies (rabbit anti-mouse IgG AlexaFluor 488 (Abcam catalog number ab150125), 1:500; goat anti-rat IgG Alexa Fluor^®^ 568 (Abcam catalog number ab175476), 1:500) and Hoechst stain (Cayman catalog number 15547, 1:1000). After three more washes in DPBS and one wash in water, coverslips were mounted using ProLong™ Gold Antifade Mountant (Thermo Fisher Scientific catalog number P10144) on a glass slide. Images were acquired on an Olympus IXplore spinning disk confocal microscope with Olympus objective 60x (UPLSAPO S2, NA 1. 3, oil) and 100x (UPLSAPO S, NA 1.35, oil).

### Cryo-EM

ABCB11 was purified similarly to the method described above, except that the construct used lacked GFP and instead contained a C-terminal 8xHis-Flag-StrepII-3C tag. The solubilized protein was batch bound to Co-NTA resin (Takara Bio USA), washed with buffer A with 10 mM imidazole, omitting DTT, and eluted using buffer A with 300 mM imidazole, omitting DTT. The remaining steps were performed as previously described. Prior to freezing, ABCB11 was concentrated to 2 mg/mL. Vitrification was performed using a Vitrobot Mark IV on Quantifoil HexAuFoil grids, applying 3.5 µL of the sample with a blot time of 5 seconds and a blot force of 10. Cryo-EM data collection was conducted on a Thermo Fisher Titan Krios operating at 300 kV at the Electron Microscopy Core Facility at UMass Chan Medical School. The microscope was equipped with a Gatan K3 direct electron detector and an energy filter set to a slit width of 20 eV. Automated data acquisition with SerialEM resulted in the collection of 17,423 movies over 24 hours, with a total dose of approximately 50 e/Å^2^. Image processing using CryoSPARC ^49^ resulted in 435,418 particles for the final reconstruction, achieving a resolution of 2.78 Å based on the FSC 0.143 criterion. Model building was initiated by fitting PDB ID 6LR0 into the density map, followed by iterative rounds of rebuilding with ISOLDE ^50^ in ChimeraX ^51^ and real-space refinement using PHENIX ^52^.

## Data availability

BSEP structural coordinates and density maps will be deposited in the PDB and EMDB upon acceptance.

## Acknowledgements

The authors would like to acknowledge Thomas Schwartz and the members of the Schwartz Lab, Department of Biology, MIT, for their scientific support. We would also like to thank the Lomason Lab, Department of Biology, MIT, for the use of their confocal microscope, as well as Jaime Cheah and Christian Soule from the MIT High-Throughput Sciences Facility for their technical expertise, training and support. We thank the Electron Microscopy Core Facility at UMass Chan Medical School, especially Christna Ouch, for their assistance.

## Author contributions

C.G. was responsible for study design and all CETSA and confocal microscopy experiments. B.R. carried out BSEP protein purification and cryo-EM structure determination. J.M. supervised the research and wrote the manuscript.

