## Supplementary information for "A structural and mechanistic model for BSEP dysfunction in PFIC2 cholestatic disease"

**Supplementary Material**

**Figure 1**

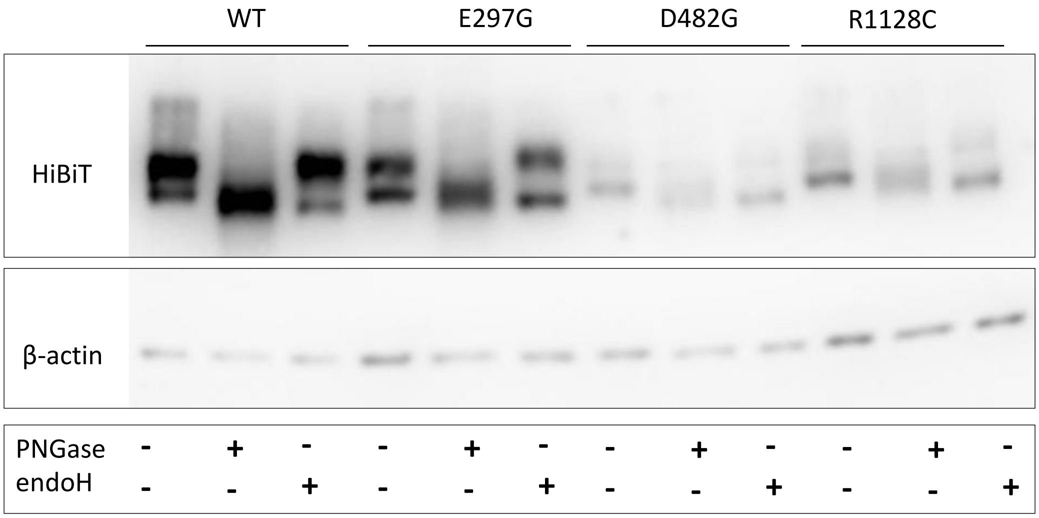


**PNGase and EndoH treatment of glycosylated BSEP proteins**

To confirm the glycosylation state of B and C-band BSEP proteins, lysates were treated with PNGase F or EndoH (New England Biolabs) in denaturing conditions following the provider's instructions. Western Blots were performed as described in Methods. For better visualization, the amount of cell lysate loaded on gel was twofold for mutants compared with WT. The mature complex glycosylated band is sensitive to PNGase F only, whereas the immature core-glycosylated band is sensitive to both PNGase F and Endo H.

**Figure 2**

**a**

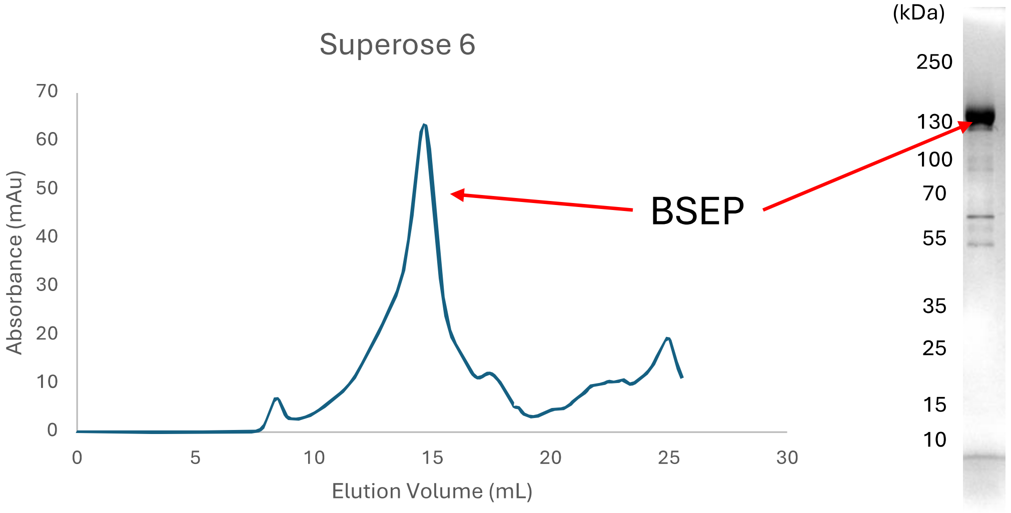


**b**

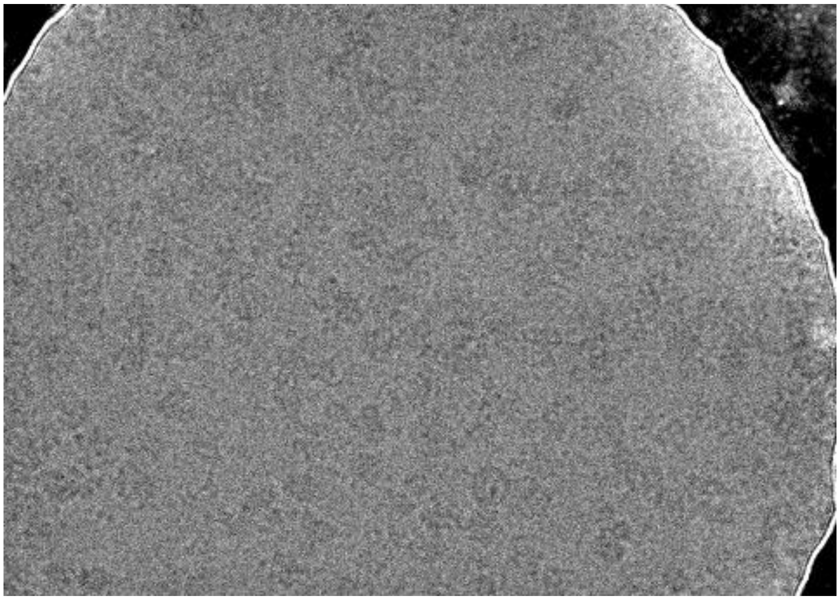

**BSEP purification and cryo-EM micrograph**

a) Size exclusion chromatography on a Superose 6 Increase column and SDS-PAGE gel of the major peak. b) A representative cryo-EM micrograph on a HexAuFoil grid imaged at 215kx.

**Figure 3**

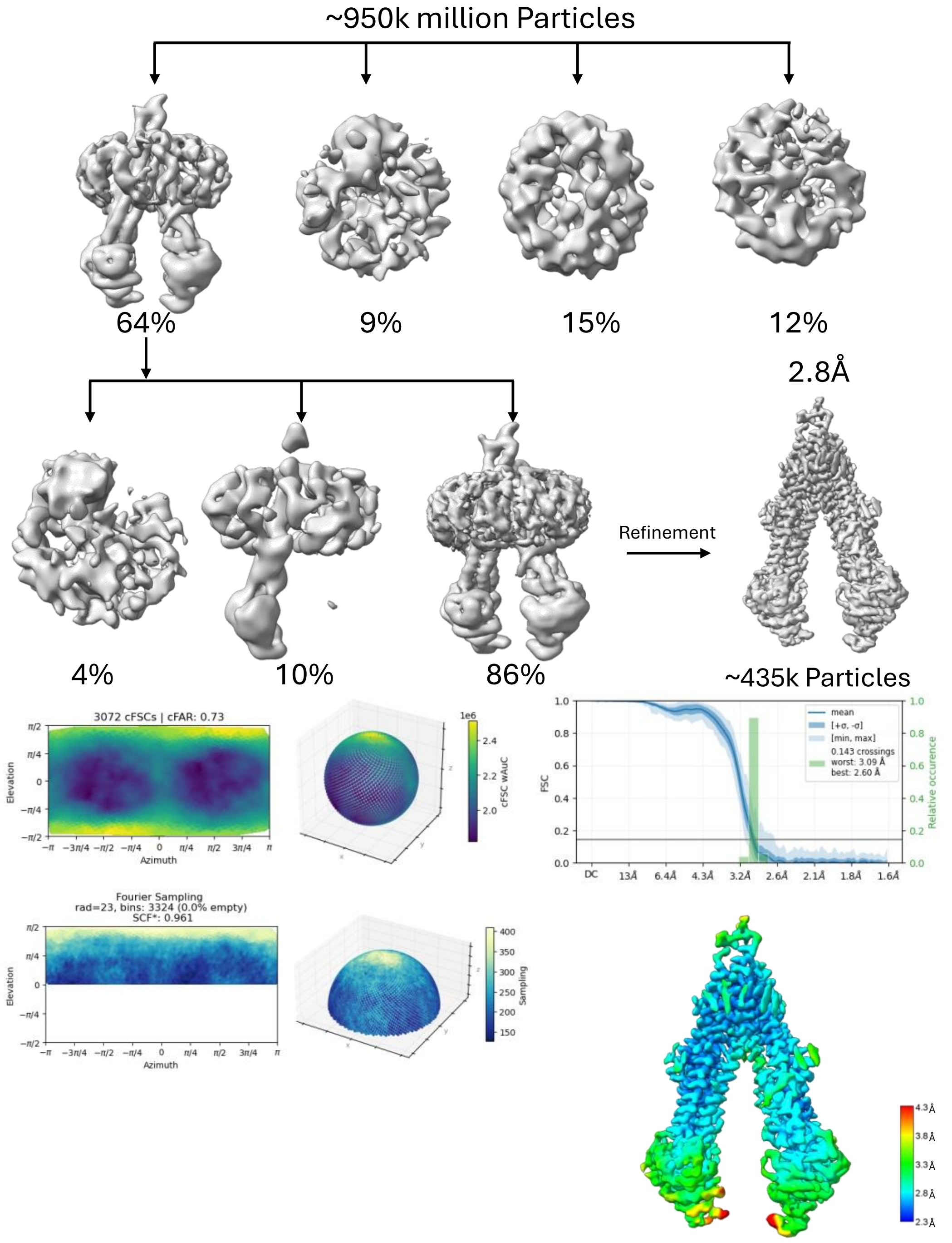


**Cryo-EM 3D-reconstruction flow-chart**

Using template picking, ~950k particles went through two rounds of heterogenous refinement with ab-initio models, followed by non-uniform refinement in CryoSPARC. The orientation diagnostics show no orientation preference issues, and the FSC curve shows the resolution estimate of 2.8Å visualized in the local resolution colored map.

**Table 1**

**Table 1: Statistics of Data Collection, Reconstruction, and Model Building**

| | |
|---|---|
| **Data Collection** | |
| EM equipment | Thermofisher Titan Krios G3 |
| Voltage (kV) | 300 |
| Detector | Gatan K3 |
| Pixel Size (Å/pixel) | 0.4012 hardware binned (0.2006 super-resolution) |
| Electron dose ($e^{-}/Å^{2}$) | 49.13 |
| Defocus Range (µm) | 1 - 2 |
| Movies | 17,423 |
| Data Collection Software | SerialEM |
| **Reconstruction** | |
| Software | CryoSPARC |
| Number of particles used | 435,418 |
| Symmetry | C1 |
| Final Resolution (Å) | 2.8 |
| **Model** | |
| Model Building Software | Coot, Isolde |
| Model Refinement Software | PHENIX |
| Map CC (mask) | 0.78 |
| Map CC (peaks) | 0.43 |
| Map CC (volume) | 0.77 |
| Rmsd (bonds) (Å) | 0.004 |
| Rmsd (angles) (º) | 0.58 |
| **Validation** | |
| MolProbity score | 1.43 |
| Clash score | 4.63 |
| Ramachandran Plot | |
| Outliers(%) | 0 |
| Allowed(%) | 3.19 |
| Favored | 96.81 |
| Rotamer outliers (%) | 0.88 |
| Cβ outiers (%) | 0 |
